# INFORME: coupling information-theoretic experimental design with nonlinear mixed-effects modeling for efficient observation scheduling

**DOI:** 10.64898/2026.08.28.747735

**Authors:** Heyrim Cho, Tingting Tang, Allison Lewis, Kathleen Storey, Tin Phan

**Affiliations:** School of Mathematical and Statistical Sciences, Arizona State University, Tempe, AZ, USA; Department of Mathematics and Statistics, San Diego State University, San Diego, CA, USA; Department of Mathematical Sciences, Lafayette College, Easton, PA, USA; Department of Medicine, University of Minnesota, Minneapolis, MN, USA; Theoretical Biology and Biophysics, Los Alamos National Laboratory, Los Alamos, NM, USA

**Author notes:** (HC); (TP).

## Abstract

Mathematical models of treatment response can inform individualized therapy, but their calibration often requires longitudinal measurements that are costly, burdensome, and collected on fixed schedules. Such schedules may be inefficient, over-sampling patients whose response is already well characterized while delaying informative measurements for those whose model parameters remain uncertain. We present INFORME (INFORmation-theoretic design with Mixed Effects), a framework that combines Bayesian information-theoretic experimental design with nonlinear mixed-effects modeling to adaptively select each patient’s next measurement time. Population and response-subgroup parameter distributions learned from an existing cohort provide informative priors, allowing candidate measurement times to be ranked by their expected reduction in patient-specific parameter uncertainty. As observations accumulate, priors can be updated to reflect the response subgroup most consistent with the patient’s data. We evaluate INFORME in two radiotherapy datasets: 150 synthetic tumor volume trajectories from a hybrid cellular automaton model of prostate cancer spheroids (HD1) and longitudinal tumor volumes from 39 patients with head-and-neck cancer (HD2). In HD1, population priors allowed omission of both pretreatment scans, while adaptive scheduling reduced the protocol from nine scans to three or four, with the response group identified from a single post-treatment scan on day 27. In HD2, the adaptive schedule used three scans instead of six and improved prediction by delaying the first on-treatment scan from week 1 to week 2, avoiding transient dynamics that produced false-positive and false-negative response projections. Across both datasets, the adaptive schedules used a mean of 2.7 scans in stead of seven and advanced completion of the patient-specific prediction by a mean of 15.5 days (95% CI, 6.7–24.3) relative to the equidistant protocol, while treatment duration remained unchanged. INFORME therefore reduces measurement burden and accelerates patient-specific prediction by concentrating observations at times that are most informative for model calibration.

**Author summary:** When a patient is treated for cancer, repeated measurements of tumor state track its response to treatment. Mathematical models can use these measurements to predict treatment response, but they need enough data to make reliable predictions. Measurements, such as imaging, are also costly, time-consuming, and burdensome for patients. In many studies, they are collected on a fixed schedule that does not incorporate the new information that has already been learned about an individual patient. We developed a method that answers when each patient’s next measurement would be most useful to inform model calibration. It first learns from previously treated patients how tumors typically respond. Then, as measurements are collected from a new patient, it identifies when another measurement would provide the most useful information about that patient’s response. We tested this approach using simulated tumors and measurements from patients with head-and-neck cancer. By concentrating scans at more informative times, our method could reach predictions earlier while using substantially fewer measurements. In the simulated study, a patient’s response group could be identified from the first scan after treatment began. Our results show that better-timed measurements can reduce scanning burden while providing useful information about treatment response sooner.

## Introduction

Despite advances in clinical trial design and computation, clinical development success rates remain low. Only 10.4% of Phase I development paths reached FDA approval in a survey of 5,820 phase transitions between 2003 and 2011 [1], and an analysis of 9,704 programs over 2011-2020 placed the overall likelihood of approval from Phase I at 7.9% [2]. Amid these concerns, mathematical modeling has emerged as a promising tool to improve clinical outcomes and reduce development risk, culminating in the establishment of a dedicated pathway for model-informed drug development (MIDD) at the FDA [3]. In particular, standard modeling approaches using dynamic models can be used to quantify drug effects, patient responses, and construct in silico cohorts [4, 5] or digital twins [6] to study optimal treatment regime and predict clinical trial outcomes [7]. These models recapitulate how participant response changes over time and reflect the continuous influence of clinical parameters. Thus, they provide a mechanistic explanation for trial outcomes, validating results beyond statistical significance and informing future applications.

Developing such models requires detailed data commensurate with their complexity, typically in the form of frequently sampled longitudinal measurements, to enable robust parameter calibration and continuous tracking of clinical endpoints [8, 9]. However, high costs, measurement errors, and variable patient compliance often limit data collection [10, 11]. Moreover, patient individual difference makes fixed sampling schedules impractical, as the fixed sample size may be too few to fully capture some patients’ response dynamics, leaving parameters underdetermined, while requiring too many samples risk noncompliance and escalating costs. An adaptive data-collection protocol that tailors sampling in real time to each patient’s response, while targeting for the data needed for model calibration, can directly address these challenges.

Two well-established bodies of work are relevant to this problem. Bayesian optimal experimental design frames measurement selection as the problem of maximizing the expected information gained about model parameters [12]. Over the past two decades, a range of computational methods have made this approach increasingly practical for nonlinear models [13, 14]. In systems biology, mutual-information criteria have been used to choose informative experiments for biochemical network models [15, 16], while more recent work has explored policy-based methods for sequential experimental design in ordinary differential equation models [17]. A separate line of work comes from population optimal design in pharmacometrics, where sampling schedules are chosen by optimizing a function of the population Fisher information matrix for nonlinear mixed-effects models [18, 19]. These methods are implemented in widely used software [20, 21] and are routinely applied in industry. Despite addressing closely related questions, the two fields have largely developed independently. Information-theoretic approaches have focused mainly on designing measurements for individual subjects under assumed parameter priors, whereas population optimal design typically produces a single sampling schedule that is fixed in advance and applied across an entire cohort. Pharmacometric researchers have identified adaptive, individual-level experimental design as an important unmet methodological need [22]. INFORME is designed to bridge this gap.

We propose coupling Adaptive Information-theoretic Design Experimentation, referred to here as AIDE, with Nonlinear Mixed-Effects modeling (NLME) in a unified INFORmation-theoretic design with Mixed Effects (INFORME) framework to optimize data-collection schedules. AIDE is a Bayesian framework that sequentially suggests the timing of each measurement to maximally reduce parameter uncertainty given accumulated knowledge [23, 24]. However, AIDE relies heavily on prior parameter distributions, so if those priors are biased or poorly specified, it may produce suboptimal sampling. To overcome this limitation, we integrate NLME to learn those distributions hierarchically from population data [25, 26] and update them within AIDE. In this work, we apply INFORME to both simulated and clinical cancer radiotherapy datasets to demonstrate its utility in adaptively optimizing data collection for model calibration and clinical trial design. In particular, the framework proposes alternative measurement schedules that require a mean of 2.7 measurements per patient instead of seven, a reduction of approximately 60%, while enabling patient-specific prediction a mean of 15.5 days earlier (95% CI, 6.7–24.3 days) than the reference equidistant protocol.

## Materials and methods

### High-fidelity data (HD1): prostate cancer treated with radiotherapy

We used a hybrid cellular automaton model to generate longitudinal data of prostate cancer dynamics under treatment with radiotherapy. The cellular automaton model [27, 28] simulates a 2-D lattice cross-section of a spheroid of heterogeneous prostate cancer cells. Each lattice site hosts one cell exposed to a spatially varying oxygen field that modulates its growth dynamics while it progresses through a stochastic cell cycle with a specified mean cell-cycle time 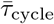. Radiotherapy is applied via the linear-quadratic model [29, 30], where the survival fraction of cells after an administered dose *d* is 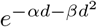 with *α* and *β* representing tissue-specific radiosensitivity parameters. The volume of the spheroid is recorded over time and used as the high-fidelity tumor-volume data.

To create a virtual cohort with diverse treatment responses, we simulated 150 tumor spheroids by systematically varying the mean cell-cycle time 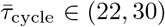 hours, the radiosensitivity parameter *α* ∈ {0.014, 0.14}, and the probability of necrotic cell being removed *ϕ* ∈ (0.005, 0.01), while fixing *α/β* = 1.5 [31] and other parameters at previously calibrated values [27, 28]. We classified the data into three response groups based on the ratio of tumor volume at the end of treatment to the peak tumor volume. Of the 150 simulated tumors, 120 were used for this classification, with 40 in silico tumors assigned to each group. High responders had a tumor volume decrease of at least 90%, medium responders had a decrease between 20% and 90%, and low responders had a decrease of less than 20% or an increase in tumor volume.

### High-fidelity data (HD2): head and neck cancer treated with radiotherapy

We used Plot Digitizer (https://plotdigitizer.com/) to extract longitudinal tumor volume measurements for 39 patients with head and neck cancer (17 from H. Lee Moffitt Cancer Center and 22 from MD Anderson Cancer Center) undergoing radiotherapy at Moffitt Cancer Center and MD Anderson Cancer Center [32]. All patients received 66-70 Gy in approximately 2 Gy weekday fractions, using either standard daily fractionation or accelerated fractionation. We assessed the temporal accuracy of the digitization by comparing the extracted treatment start time with its known value of day 0. The mean error was 0.04 days (range: 0–0.10 days), indicating good agreement with the known treatment start time. Because no reference values were available for the remaining time points and tumor volume, their digitization accuracy could not be assessed directly. We therefore also visually inspected the extracted time series for obvious inconsistencies.

For these digitized patient data, we classified individuals into three response groups based on the ratio of tumor volume at the last measured time point to the tumor volume immediately before treatment (time 0). High responders had a ratio below 1/3, medium responders had a ratio between 1/3 and 2/3, and low responders had a ratio above 2/3. Based on this classification, there were 14 high responders, 14 medium responders, and 11 low responders.

### A mathematical model of prostate cancer response to radiotherapy

The logistic growth model forms the backbone of many cancer models, including prostate cancer [33, 34]. Cho et al. previously demonstrated the utility of AIDE to optimize data collection design for prostate cancer under radiotherapy [24]. Thus, we also used a simple logistic model as our low-fidelity model for HD1 dataset. Let *x*(*t*) tracks the tumor volume, then its growth dynamics follows

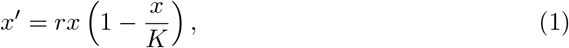

where *r* and *K* are the intrinsic growth rate and carrying capacity, respectively. We track the radiotherapy dose and effect via *R*(*t*) and *γ*(*t*), respectively. At the time *t*_*i*_ of each treatment, we model the radiation dose *R*(*t*) delivered at rate *f* (*t*_*i*_), where 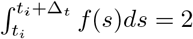 Gy with Δ_*t*_ being the duration of each radiotherapy treatment. The radiation decays rapidly at rate *k*, fixed at a large value of 100 per day, or about 4.17 per hour, so that the delivered dose is effectively instantaneous on the time scale of tumor growth. The radiation reduces tumor volume at constant rate *γ*_0_ (day^−1^), only if *R*(*t*) *>* 0.001 Gy. This formulation approximates the constant reduction of tumor volume during each session of radiotherapy. The full model is

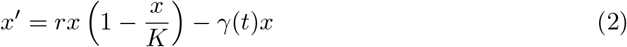

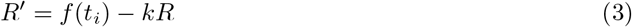

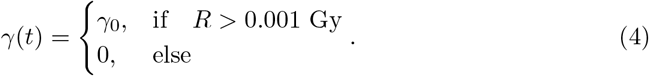

### A mathematical model of head and neck cancer response to radiotherapy

The HD2 data set was previously studied with several mathematical models by Zahid et al. and Mohsin et al. [32, 35]. In the model of Zahid et al., the carrying capacity is reduced by a fixed fraction at each fraction of radiotherapy, *K*_+_ = *K*_−_(1 − *δ*), where *K*_−_ and *K*_+_ denote the carrying capacity immediately before and after a fraction, respectively. Here, we use a continuous-time variant of that mechanism, which allows the carrying capacity to be evaluated at arbitrary candidate measurement times as required by the design algorithm, and which follows the notation used for HD1:

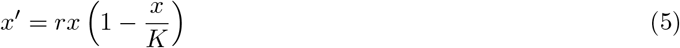

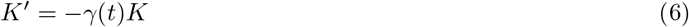

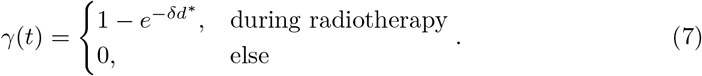

In this model variant, radiotherapy reduces the carrying capacity *K* at a rate 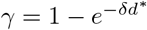 only during the radiotherapy course. *δ* is the radiation sensitivity parameter, and *d*^∗^ = *d/*Δ_*t*_ is the dose rate. We set *x*(0) to the initial tumor volume measurement for each individual, while *K*(0) is the ratio of the initial volume with the tumor proliferative saturation index (PSI), *K*(0) = *x*(0)*/*PSI [36].

## INFORME

**In the information theoretic design framework**, we consider a high-fidelity model **ℳ**_***H***_ (***θ***, *t*_*n*_) as a realistic, computational or experimental, models used for data generation, while a low-fidelity model **ℳ**_***L***_(***θ***, *t*_*n*_) as a coarse-grained computational model used for data fitting and downstream analysis. ***x***(*t*) ∈ ℝ^*K*^ is the *K*-dimensional state vector, ***θ*** ∈ ℝ^*M*^ is the *M*-dimensional parameter vector, and *t*_*n*_ is the discrete time corresponding to the data 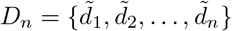, where 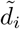 is the observation at time *t*_*i*_. According to Bayes’ rule, the posterior distribution of the parameters ***θ*** given data *D*_*n*_ is:

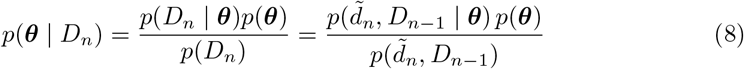

where *p*(*D*_*n*_ | ***θ***) is the likelihood of obtaining data *D*_*n*_ given parameter set ***θ***, and *p*(***θ***) is the prior distribution of ***θ***.

The mutual information between the parameter vector and a future observation is defined as:

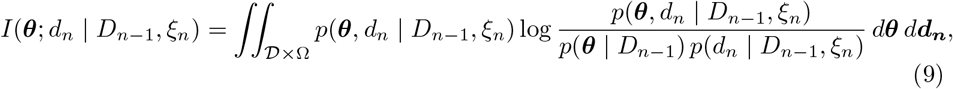

where Ξ = {*ξ*_1_, …, *ξ*_*N*_*}* is the set of all admissible designs, with *t*_*N*_ the final candidate time, *d*_*n*_ is the predicted value of 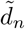 using a low-fidelity model, *D* is the complete set of all (unknown) future observations, and Ω ⊆ ℝ^*M*^ is the admissible parameter space.

The mutual information quantifies the expected uncertainty reduction in the estimation of ***θ*** by taking the next measurement *d*_*n*_ at time *t*_*n*_ according to design *ξ*_*n*_. For example, suppose we have a data set *D*_*n*−1_ consisting of high-fidelity measurements *{d*_1_, …, *d*_*n*−1_*}* and a set of all possible designs Ξ. For each candidate design *ξ*_*n*_ ∈ Ξ, we consider the hypothetical addition of a new high-fidelity data point *d*_*n*_ obtained under design *ξ*_*n*_ to the existing data set *D*_*n*−1_. The design *ξ*_*n*_ that maximizes the mutual information is the one that reduces the most uncertainty in the estimation of ***θ***.

A limitation of the canonical mutual information is that it may favor an optimal design time point far in the future, creating an excessively large gap between *d*_*n*−1_ and *d*_*n*_. In clinical settings, this could translate into substantial treatment delays, added resource use, or missed opportunities to test alternative therapies. To address this, we adopt the adaptive score function of Cho et al. [24], which penalizes designs with excessively long waiting times to reflect the information lost by skipping the intermediate candidate times.

Let *r* denote the index of the most recently collected measurement, so that 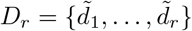 is the current data set and 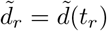 is the most recently appended data point. The remaining candidate designs are *ξ*_*r*+1_, …, *ξ*_*N*_. For each of these we evaluate the mutual information *I*(***θ***; *d*_*i*_ | *D*_*r*_, *ξ*_*i*_) and rescale it to the unit interval,

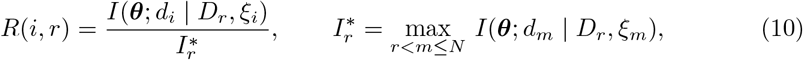

so that *R*(*i, r*) ∈ [0, 1], with equality to one only for the design that maximizes the mutual information. The score function for candidate design *i* at step *r* is then

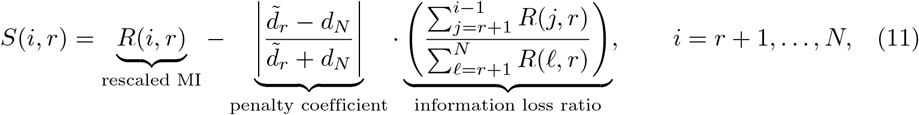

and the next measurement is taken at the design 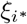, where *i*^∗^ = arg max_*i*_ *S*(*i, r*).

The penalty coefficient is a symmetric absolute error comparing 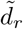, the high-fidelity measurement, with *d*_*N*_, the anticipated final measurement given by the low-fidelity model prediction for the final day of treatment (both quantities are available at the time the design decision is made). It is bounded on [0, 1] and is set to zero when 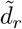 and *d*_*N*_ are both zero. A quantity still expected to change substantially over the remaining treatment period therefore incurs a large penalty for skipping data points, whereas one that has stabilized near its anticipated final value incurs a penalty close to zero, allowing for sparser data collection. The information loss ratio totals the rescaled mutual information of the points *r* + 1 through *i* − 1 that would be skipped by measuring at *t*_*i*_, divided by the sum across all remaining candidate designs, so that the penalty grows with the length of the proposed gap.

The posterior distribution in Eq. (8) is computed using Markov Chain Monte Carlo (MCMC) sampling, implemented via the Delayed Rejection Adaptive Metropolis (DRAM) algorithm [37]. For each calibration, the chains are simulated for an initial 2,000 iterations discarded as a burn-in period to ensure chain convergence, followed by 10,000 iterations to characterize the parameter posterior distributions. The mutual information in Eq. (9) involves a high-dimensional integral and this is typically estimated via the kth-nearest neighbor (kNN) method [38].

**In the nonlinear mixed-effects modeling framework**, the parameters in the low-fidelity model are treated as random effects representing a population distribution. We denote 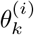 the *k*-th parameter for the i-th individual, and 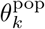 the respective mean for the population. Then, assuming a log-normal distribution, the relationship between individual and population parameters is

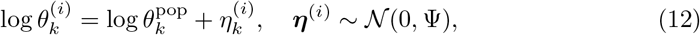

where ***η***^(*i*)^ is the vector of random effects (inter-individual variability) for individual *I* and Ψ is its covariance matrix. Monolix reports 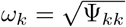, the standard deviation of the *k*-th random effect; these are the quantities given in Tables 2 and 4. The low-fidelity model and its connection with the data generated from the high-fidelity model is:

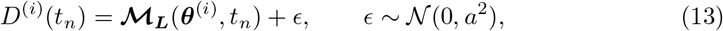

where *D*^(*i*)^(*t*_*n*_) is the data generated for individual *i* and *ϵ* is the measurement error, taken as additive with standard deviation *a* estimated jointly with the population parameters.

NLME analysis is performed using the software Monolix (Lixoft), which estimates population parameters by the stochastic approximation expectation maximization (SAEM) algorithm [39]. Candidate models are compared using the −2 log-likelihood (−2LL), estimated by importance sampling, and the corrected Bayesian information criterion (BICc) [40], both reported by Monolix.

### How NLME Fits into the MI Framework

In practice, we can usually only use the data generated from the high-fidelity model without full knowledge of all of the biological parameters used in the high-fidelity model to generate it. Therefore, we seek a reasonable low-fidelity model to approximate the high-fidelity model based on the high-fidelity generated data. Instead of assuming an arbitrary prior in the MI framework, NLME allows us to estimate the population distribution of each parameter, which we can then use as a prior in the MI framework. Taking this one step further, we can characterize sub-population priors based on response to treatment by incorporating a covariate on key parameter affecting the response. For example, if *θ*_*x*_ is a key parameter associate with treatment response, then we can characterize the priors for the strong and weak responders by incorporating a covariate to *θ*_*x*_ in the data fitting.

When initiating the INFORME algorithm, we start with a general population distribution for the parameters (i.e., no distinction based on response to treatment). As new measurements become available, we can iteratively update the distribution of parameters using the distinct sub-population distributions. Incorporating this additional prior information can help the algorithm converge faster in some scenarios. The iterative procedure continues until the absolute change |*θ*_*n*_ − *θ*_*n*−1_| and the relative change |*θ*_*n*_ − *θ*_*n*−1_|*/θ*_*n*_ in the posterior mode of the treatment-related parameter (*γ*_0_ for HD1, *δ* for HD2) between successive measurements both fall below prescribed tolerances. Convergence is assessed on the stability of the posterior mode instead of the width of the credible region because the credible region might be underestimated when only a small number of observations is available. If neither tolerance is met, the procedure terminates when the candidate set is exhausted at the end of the treatment course, so that an adaptive schedule never requires more measurements than the equidistant protocol. Because these criteria are evaluated for each individual, the resulting schedules differ across patients. For presentation and group-level comparisons, each response group is summarized by the modal scan time at each step across individuals in that group. Collapsing the individualized schedules into a single fixed schedule per group provides a conservative estimate of the benefit of fully individualized scheduling. A schematic of the workflow is illustrated in Fig 1.

**Fig 1.**
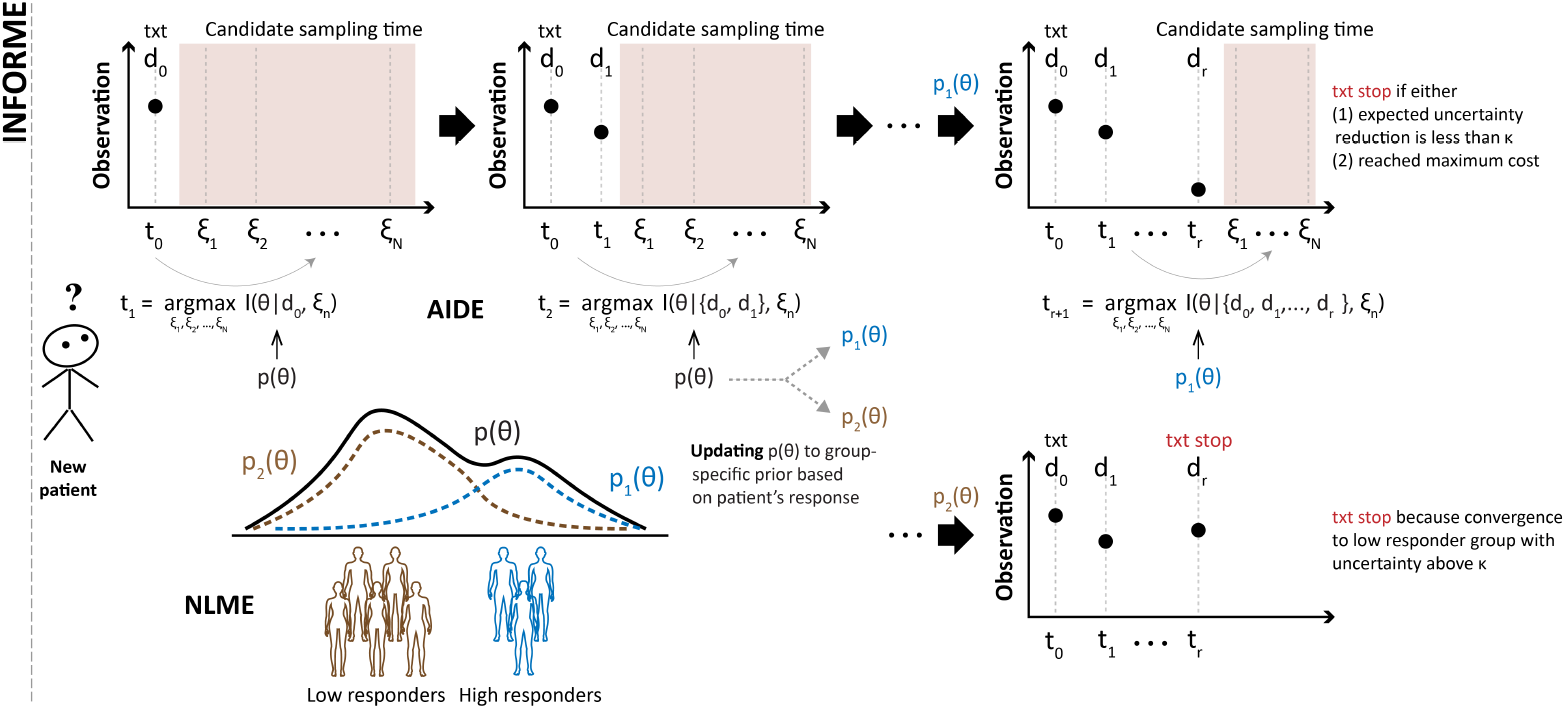
Schematic of the INFORME framework. A new patient enters the clinic with no individual measurement. A population-level parameter prior *p*(***θ***), estimated by NLME from a previously calibrated cohort, provides the starting point (lower row). At each step the design criterion evaluates the admissible candidate measurement times *ξ*_1_, …, *ξ*_*N*_ and selects the time maximizing the score of Eq (11) (upper row, AIDE). After some post-treatment measurements, the individual is assigned to a response group and the population prior is replaced by the corresponding group-specific distribution *p*_1_(***θ***) or *p*_2_(***θ***) estimated from the response subpopulations (lower row, NLME). Subsequent measurement times are selected under the updated prior. The procedure stops when the expected uncertainty reduction falls below a threshold *κ*, when the parameter estimates have converged, or when the measurement budget or the end of the treatment window is reached. Two response groups are shown for clarity with *p*_1_(***θ***) and *p*_2_(***θ***) denoting the high- and low-responder priors. *κ* denotes the tolerance on the expected uncertainty reduction below which no further measurement improves model prediction.

### Evaluation metrics

We compare schedules using: (1) the *measurement burden* is the number of scans required per patient, counted from treatment initiation and including any pretreatment scan needed to initialize the model; and (2) the *time to complete prediction* is the day of the last scan that a schedule requires. For each response group, the *time saved* by an adaptive schedule is defined as the difference between the final scan day of the reference equidistant protocol and that of the adaptive schedule. Because the adaptive procedure ends once the treatment-related parameter estimate has converged, this difference represents the interval by which the patient-specific prediction is finalized earlier. It does not alter the actual radiotherapy course.

## Results

### Constructing response-based priors

We first calibrated each low-fidelity model to its cohort using NLME, and used the resulting parameter distributions as priors for the design procedure. For HD1, the model of Eqs (2)-(4) was fitted to the 120 classified virtual tumors with log-normal random effects on the intrinsic growth rate *r*, the carrying capacity *K*, and the treatment parameter *γ*_0_. For HD2, the model of Eqs (5)-(7) was fitted to the 39 patient trajectories with log-normal random effects on PSI and the radiation sensitivity *δ*. In both cases the response group entered as a categorical covariate on the treatment parameter (*γ*_0_ for HD1, *δ* for HD2), so that the growth-related parameters share a single population distribution across groups while the treatment response remains group-specific. Full population estimates are given in Tables 2 and 4.

Including the response-group covariate reduces response-group variability and subsequently allows faster convergence of the INFORME framework. In HD1, the inter-individual standard deviation of *γ*_0_ falls from 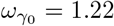 to 0.41 once the covariate is included, while the estimates of *r* and *K* are essentially unchanged (*r* = 0.32, *ω*_*r*_ from 0.12 to 0.13; *K* = 0.57, *ω*_*K*_ from 0.13 to 0.15). The group-specific typical values of *γ*_0_ are 4.12, 1.81, and 0.38 for high, medium, and low responders, spanning about an order of magnitude. The same pattern holds in HD2, where the group-specific values of *δ* differ by about eight-fold across the three groups. Because most of the between-individual variability in treatment response is absorbed by group, the group-conditional distributions are substantially narrower than the pooled population distribution, and it allows post-treatment measurements to quickly select which group an individual belongs to. These priors are constructed from an already calibrated cohort. In prospective use that cohort would consist of the previously treated patients available when a new patient enters.

### Scan number reduction from using informed priors in dataset HD1

We first compare the proposed adaptive scan schedules with other baseline schedules using the synthetic dataset HD1. The evaluated scanning protocols depend on whether they require pretreatment scans and whether they rely on equidistant versus adaptive schedule. The three scan schedules were defined as follows:

- Schedule A: Baseline equidistant schedule. This schedule includes two pretreatment scans, one scan at treatment initiation on day 15, and six subsequent weekly scans collected every Friday, for a total of nine scans.
- Schedule B: Population-informed equidistant schedule. This schedule eliminates the two pretreatment scans by using NLME-derived population priors to inform the non-treatment parameters. It begins with one scan at treatment initiation, followed by six weekly scans on a fixed weekday, for a total of seven scans. We evaluated Tuesday, Wednesday, Thursday, and Friday schedules; the Friday schedule is shown in the figure.
- Schedule C: Population- and group-informed adaptive schedule. This schedule also eliminates pretreatment scans by using NLME-derived population priors. After the scan at treatment initiation, the timing of subsequent scans is selected adaptively using mutual information. The first post-treatment scan is scheduled on day 27 based on the population prior for the treatment-related parameters. The day 27 measurement is used to classify the patient as a high (Group 1), medium (Group 2), or low (Group 3) responder. The treatment-parameter priors are updated to the corresponding group-specific distributions, and the remaining scan times are selected accordingly. Depending on the response group, Schedule C requires only two or three additional scans. The representative group-specific schedules were defined using the modal scan times across the HD1 cohort.

Figure 2 shows the example fits in each patient groups using the three scan schedules. The population and group-specific prior distributions are shown as well. Schedules B and C provide robust prediction on the radiotherapy outcome without requiring the pre-treatment scans mandated in schedule A. In addition, schedule C yields robust predictions using significantly fewer scans than the equidistanced schedules, while simultaneously enabling early prediction at mid-treatment.

**Fig 2.**
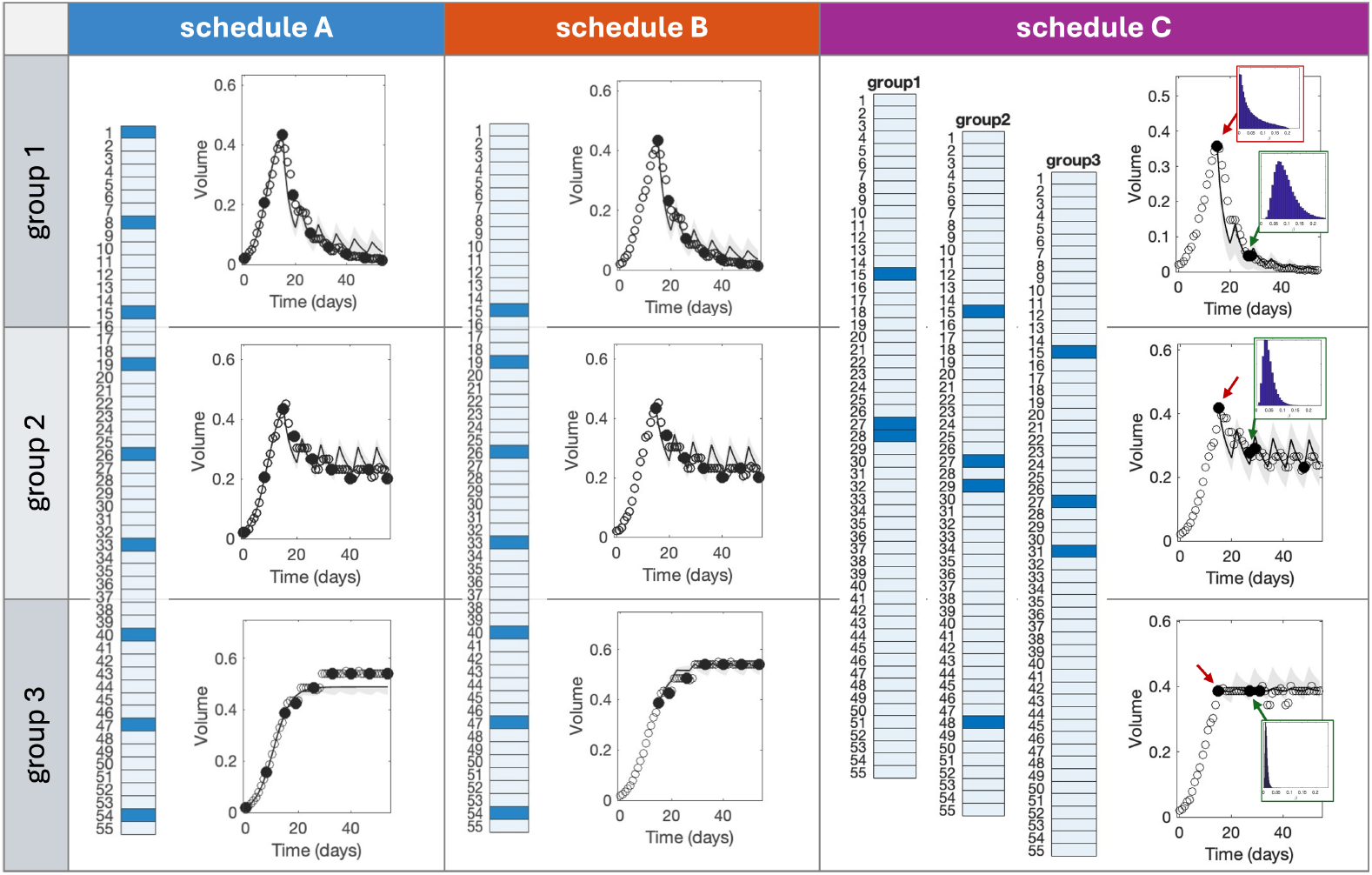
Comparison of equidistant and adaptive scan schedules and representative model fits for the HD1 dataset. Rows correspond to the three treatment-response groups: high responders (Group 1), medium responders (Group 2), and low responders (Group 3). Blue cells indicate selected scan days. Schedule A is the baseline protocol with pretreatment and weekly scans. Schedule B eliminates pretreatment scans using an NLME-derived population prior but retains weekly sampling. Schedule C uses population- and group-specific priors: the Day 27 scan classifies response, updates the treatment-parameter prior that guides subsequent scan timing. In the model-fit panels, open circles denote high-fidelity tumor-volume data, filled circles indicate observations used for calibration, solid curves show model predictions, and shaded regions represent 95% credible region area.

### Accuracy and uncertainty of adaptive scan schedule in dataset HD1

Figure 3 compares the accuracy and uncertainty of the three scanning protocols plotted against both time and total scan number. The error is computed as the mean square error between the high-fidelity data and the low-fidelity model fit during the treatment period. In Group 1 (high responders), schedule C achieves better accuracy with only two scans compared to schedules A and B that requires more scanning. The initial two scans from the equidistance schedules do not provide accurate prediction compared to the strategy of schedule C of delaying the initial scan until day 27. This phenomena is similarly observed in Group 2 (medium responders), where schedule C yields better accuracy after the initial scan. Conversely, in Group 3 (low responders), the prediction error at day 27 computed by schedule C is larger than that computed by schedule B. However, the error magnitude of using initial scan of schedule C is similar to those achieved in the other responders groups 1 and 2 that is sufficient to make robust prediction that the therapy outcome is not successful, and the accuracy significantly improves as the second scan is obtained, that eventually makes schedule C more accurate than schedule B by day 33. Moreover, the shared regions in the error versus time plots show the range of minimum and maximum errors across the patient cohort. We note that the maximum error bound under schedule C remains consistently lower than other schedules, highlighting its robust predictive performance.

**Fig 3.**
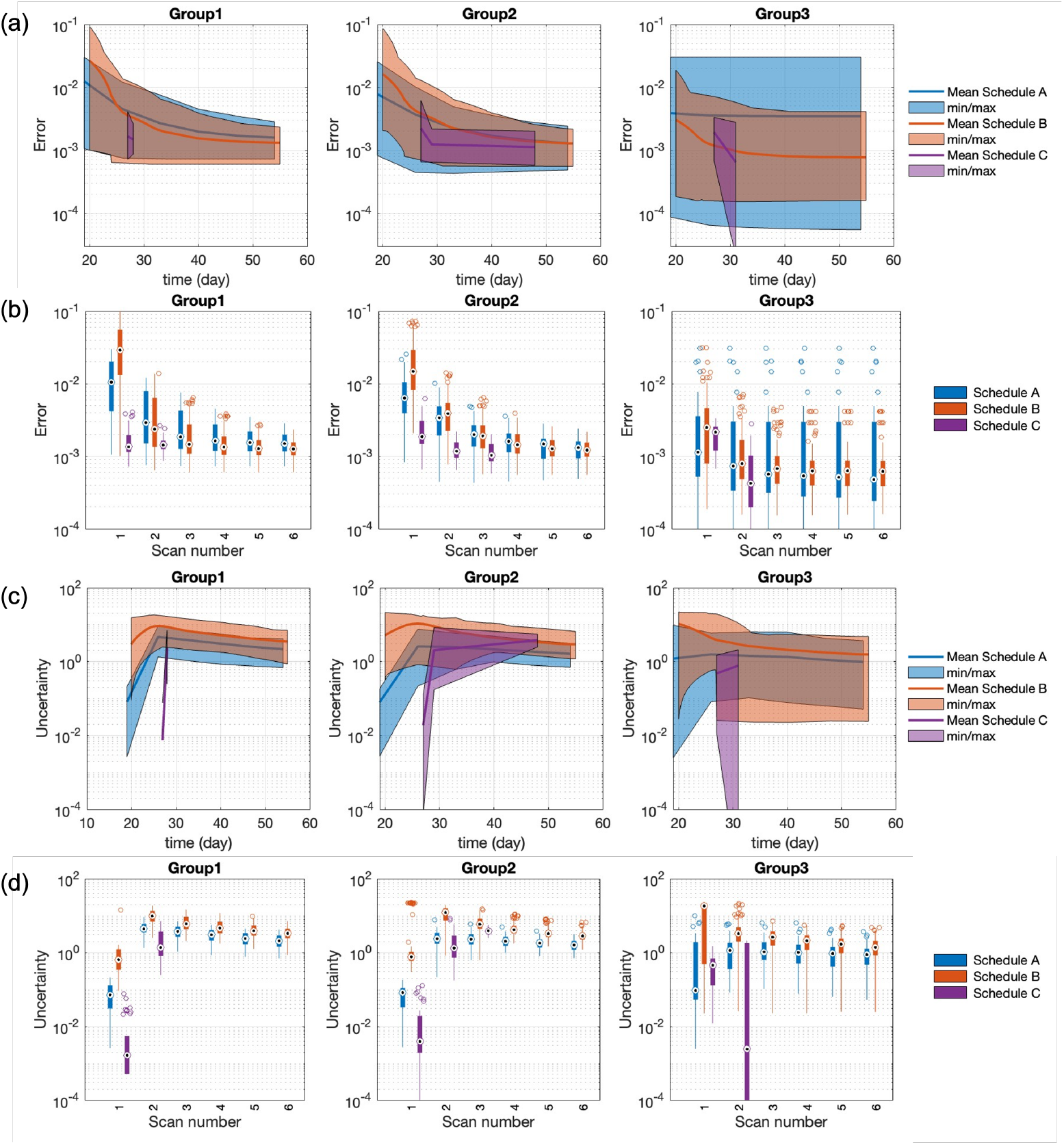
Comparison of error (a-b) and uncertainty (c-d) versus time (a,c) and scan number (b,d) across equidistant and adaptive scan schedules using HD1 dataset. The colored area in the error versus time plots is the minimum to maximum errors. The box plot shows the median, and the 25th and 75th percentiles. For high-responder group (Group 1), the selected scan schedule gives more accurate result than the equidistant schedule with fewer scans. For low-responder group (Group 3), the selected scan schedule gives similar result to equidistant schedule with fewer outlying errors.

Figure 3 also presents the uncertainty between the three schedules, where uncertainty is computed as the 95% credible region area of the model fit using the parameter posterior distribution. Uncertainty of schedule C is comparable to schedules A and B across all response groups. However, we remark that the uncertainty metric is estimated to be lower at the initial scan of schedule C at day 27. Since the model calibration relies on only two available data points at this point, the parameter space is artificially constrained, resulting in an underestimation of the predictive uncertainty. Consequently, these early-stage prediction must be interpreted with caution and should not be viewed as highly reliable predictions.

### Efficacy of adaptive scan schedule in dataset HD2

The adaptive scanning framework is also applied on the clinical dataset HD2, as shown in Fig. 4. We remark that since this dataset consists of sparse weekly measurements, the candidate decision space is far more restricted than in the synthetic dataset HD1. Furthermore, although we regard them as weekly, the exact measurement days varied across individual patients. Nevertheless, the primary advantage of the adaptive schedule is due to skipping the first week scan. Using the population prior, our proposed method prescribes initiating the scan during the second week following treatment onset.

**Fig 4.**
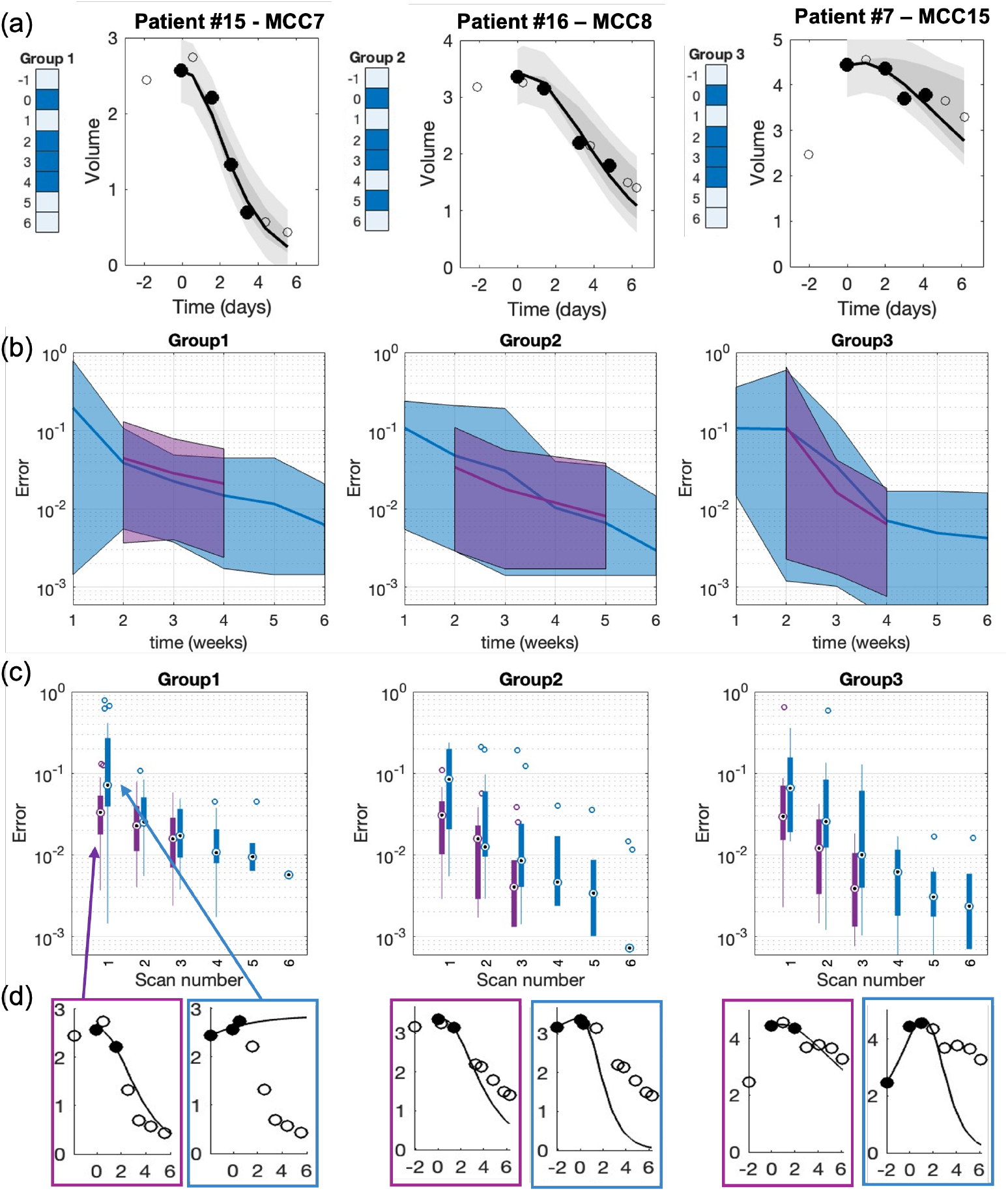
Adaptive scan schedules and model fit examples (a), error versus time (b), error versus scan number (c), and model fits after the first scan (d) for the HD2 dataset. The results compare the equidistant (blue) and adaptive (purple) schedules. By skipping the Week 1 scan, the adaptive schedule is more accurate than the equidistant schedule at each scan. Panels correspond to the three treatment-response groups: high responders (Group 1), medium responders (Group 2), and low responders (Group 3).

Compared to the initial Week 1 scan of equidistant schedule, the adaptive schedule achieves better accuracy after a single observation. This improvement is primarily driven by mitigating the impact of transient dynamics that cause Week 1 scans to yield false-positive or false-negative response projections (Fig. 4(d)). For instance, a sharp temporary increase during Week 1 for patient MCC7 results in a false-negative prediction, whereas patients MCC8 and MCC15 produces false-positives. Across all three response groups, delaying the first scan to Week 2 yields a substantially more reliable projection of the final outcome.

Following this initial observation, the adaptive framework proceeds according to group-specific schedules computed from the updated cohort priors—specifically, Weeks 2, 3, 4 for Group 1, 2, 3, 5 for Group 2, and 2, 3, 4 for Group 3. While the error with respect to scan (Fig. 4(c)) is lower under the adaptive protocol, the error evaluated over time (Fig. 4(b)) remains comparable to the equidistant baseline. Because dataset HD2 offered a very limited candidate pool (a maximum of six time points) with inter-patient variability in collection dates, there was less latitude for the adaptive schedule to demonstrate dramatic temporal error reductions over the equidistant schedule. Nevertheless, despite these clinical constraints and HD2 being substantially noisier than HD1, the adaptive scheduling framework achieves robust predictive accuracy using fewer scans than the baseline protocol.

### Group-level, population-level, and uninformative priors

We also examine the advantage of incorporating population-level and group-level prior in the adaptive framework. Specifically, we compared three prior distribution strategies which our adaptive framework:

1. Population- and group-level log-normal prior (schedule C): The initial population log-normal prior is updated to a group-specific log-normal prior following the first scan;
2. Population-level log-normal prior: The log-normal population prior remains un-updated throughout treatment; and
3. Uniform prior: An uninformative prior that provides no parameter information beyond admissibility ranges.

Fig. 5(a) shows the score function values, *S*(*i, r*) from Eq. (11), computed under both the log-normal population prior and uniform prior across HD1 and HD2 datasets. Using the population prior, the maximum score value is achieved at day 27 for HD1 and at week 2 for HD2, which was the basis of the initial scan timing chosen in schedule C. However, the uninformative uniform prior yields the score function to peak earlier, at day 22 for HD1 and Week 1 for HD2. Evaluating these three prior strategies on the HD1 dataset demonstrates that integrating group-specific updates reduces the required scanning frequency. While the uniform and static population priors both require three scans to reach convergence, Schedule C (utilizing group-specific updates) converges after two scans for Groups 1 and 3, though Group 2 still requires three. Furthermore, Schedule C achieves better predictive accuracy at the last measurement compared to both the uniform and population-level priors. These findings demonstrate that informing the adaptive framework with structured, cohort-derived prior distributions enhances prognostic accuracy while further reducing the scanning burden. We note, however, that this advantage is modest; reducing the requirement from three scans to two represents an incremental gain, and the final prediction errors remain within the same order of magnitude.

**Fig 5.**
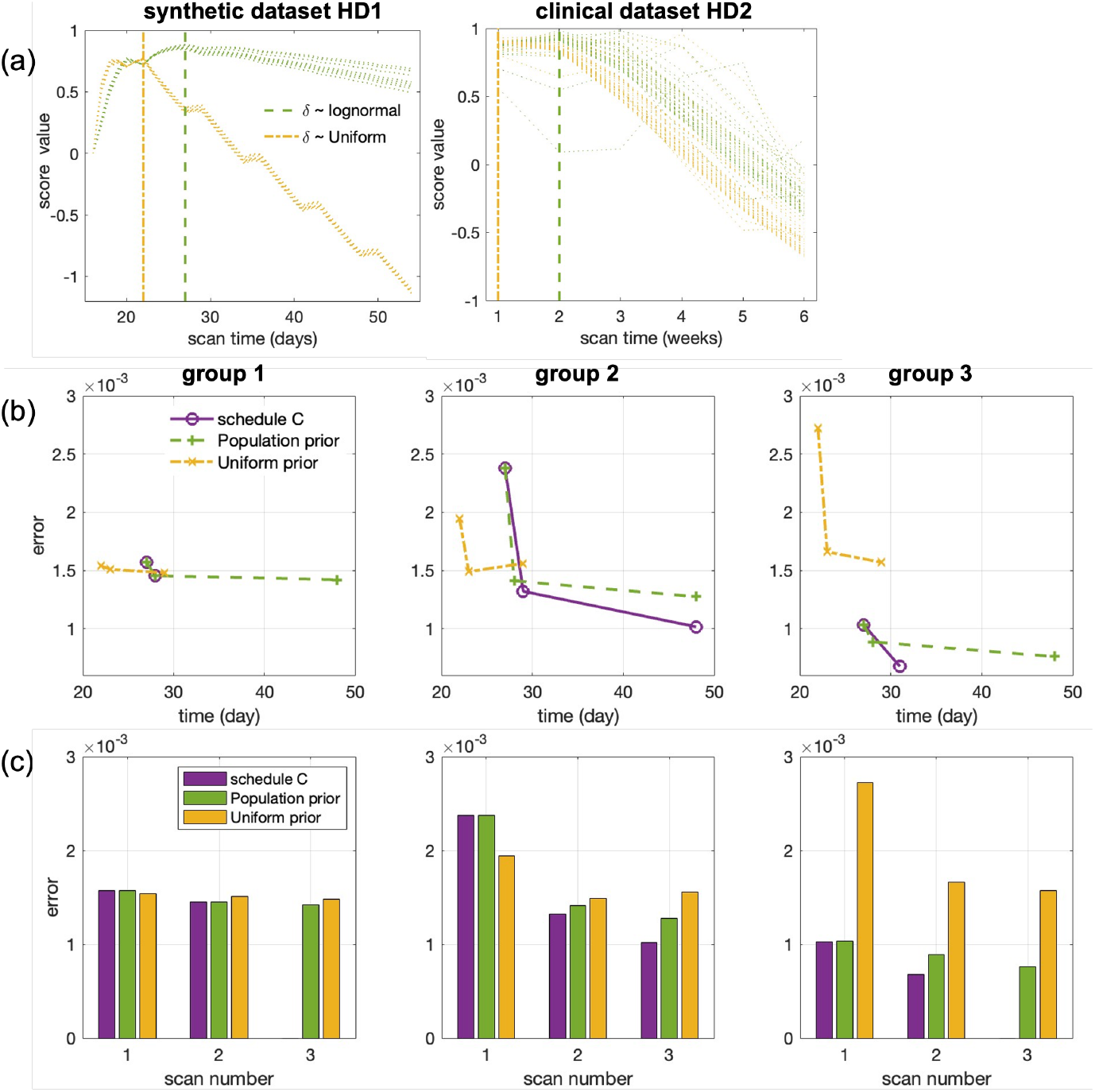
Comparison of adaptive scanning schedules under different prior distributions: uniform, population-level log-normal, and group-level log-normal. (a) Optimal initial scan timing based on maximum score values, demonstrating that population log-normal priors prescribe an initial scan during the second week across HD1 and HD2, whereas uniform priors recommend earlier initial imaging. Model error evaluated over (b) time and (c) total scan count, illustrating that schedule C informed by both population and group priors achieves better predictive accuracy with fewer total scans. In (a), the legend entry *δ* denotes the treatment-related parameter of each dataset: *γ*_0_ for HD1 and *δ* for HD2. In (b) and (c), all three conditions use the adaptive schedule and differ only in the prior. “Schedule C” denotes the population- and group-informed prior. Panels correspond to the three treatment-response groups: high responders (Group 1), medium responders (Group 2), and low responders (Group 3).

### Reduction in measurement burden and time to completed prediction

Table 1 summarizes measurement burden and prediction completion time across both datasets. In HD1, the final adaptive scan occurs on days 28, 31, and 48 for the high-, medium-, and low-response groups, respectively, compared with day 55 under the reference protocol. These schedules complete the patient-specific prediction 27, 24, and 7 days earlier, for a mean saving of 19.3 days. In HD2, the final adaptive scan occurs at Week 4 for Groups 1 and 3 and Week 5 for Group 2, compared with Week 6 under the reference protocol, yielding savings of 2, 1, and 2 weeks, respectively (mean 1.7 weeks). Across all six group-level comparisons, prediction is completed a mean of 15.5 days earlier than under the reference protocol (SD 8.4 days; 95% CI 6.7–24.3 days).

**Table 1.** Measurement burden and time to completed prediction. The reference protocol comprises one pretreatment scan and six weekly on-treatment scans (seven scans total). Adaptive scans are counted from treatment initiation. Time saved is the difference between the last scan of the reference protocol and that of the adaptive schedule.

| Dataset | Group | Last scan<br>Ref./adaptive | Time saved (d) | Adaptive scans |
| --- | --- | --- | --- | --- |
| HD1 | 1 (high) | day 55 / day 28 | 27 | 2 |
| HD1 | 2 (medium) | day 55 / day 31 | 24 | 3 |
| HD1 | 3 (low) | day 55 / day 48 | 7 | 2 |
| HD2 | 1 (high) | week 6 / week 4 | 14 | 3 |
| HD2 | 2 (medium) | week 6 / week 5 | 7 | 3 |
| HD2 | 3 (low) | week 6 / week 4 | 14 | 3 |
| **Mean, both datasets** |  |  | **15.5** | **2.7** |
| 95% confidence interval |  |  | 6.7–24.3 | — |

The adaptive schedules require a mean of 2.7 scans per patient, compared with seven under the reference protocol, which consists of one pretreatment scan and six weekly on-treatment scans. This corresponds to an approximately 60% reduction in measurement burden. In HD1, the time saving is greatest for high responders, whose trajectories separate from the population distribution early, and smallest for low responders, whose limited change in tumor volume leaves the treatment-related parameter poorly determined until later in treatment. In HD2, the restriction to six weekly candidate slots reduces the variation among groups.

## Discussion

In this work, we present INFORME, a framework for adaptive, patient-specific clinical data collection that combines Bayesian information-theoretic experimental design with nonlinear mixed-effects modeling. The framework first learns population- and subgroup-specific parameter distributions from existing cohorts, then sequentially selects future measurement times that maximize expected information gain while accounting for the cost of delaying observations. Unlike conventional fixed sampling schedules, INFORME adapts data collection to each patient’s evolving response by updating parameter priors as new measurements become available. We demonstrate the framework using both a high-fidelity synthetic radiotherapy dataset (HD1) and a clinical head-and-neck cancer dataset (HD2). Across both datasets, INFORME reduces the number of required scans while maintaining predictive accuracy and enabling earlier identification of treatment response compared to equidistant schedule. These results illustrate that integrating population-level prior information with information-theoretic adaptive scheduling provides an effective strategy for improving model calibration, reducing the number of required patient measurements, and increasing the efficiency of longitudinal clinical studies.

INFORME combines ideas from Bayesian optimal experimental design [**?**, 12–14] and population optimal design in pharmacometrics [18–21]. It uses expected reduction in parameter uncertainty to rank candidate measurement times and updates those rankings sequentially as data accumulate [23, 24]. In addition, it uses a hierarchical model of between-patient variability, but treats the fitted population and subgroup distributions as patient-specific priors rather than using them to optimize a single fixed cohort schedule. In our demonstration, these priors were informative enough to eliminate pretreatment imaging, while the distinct subgroup distributions allowed a single post-treatment measurement to assign patients to a response group and adapt the timing of subsequent scans. The contribution is a merge of established frameworks [24, 39, 40] in a sequential, individual-level design and the demonstration that cohort-derived information can substantially reduce imaging burden in real clinical data.

Further validation is needed to establish the robustness and generalizability of our framework. The present study evaluates the framework using only two radiotherapy datasets: a high-fidelity synthetic dataset and a clinical head-and-neck cancer dataset with sparse, approximately weekly measurements. In the clinical dataset, the limited number and irregular timing of available observations substantially constrained the candidate scheduling space, reducing the opportunity to evaluate the full benefit of adaptive measurement selection. Future studies should therefore examine this framework using preclinical datasets with denser longitudinal sampling, more feasible candidate measurement times, and controlled acquisition conditions. Such data would enable more rigorous comparisons between adaptive and fixed schedules, assessment of sensitivity to measurement noise, patient variability, prior mis-specification, and model discrepancy. This additional validation will be important for determining whether the observed reductions in measurement and improvements in early prediction persist across experimental systems before the framework is translated into prospective clinical studies.

The performance of INFORME may also be sensitive to several user-specified and data-dependent choices, including the convergence or stopping threshold, the assumed prior distributions, the definition of response subgroups, the level and structure of measurement noise, and the mathematical model used for prediction. In the present study, the population- and group-informed priors were evaluated primarily through updates to treatment-related parameters, and the robustness of the selected schedules to uncertainty in other biological parameters was not systematically examined. Moreover, each dataset was analyzed using a single selected mechanistic model, so the extent to which the recommended scan times depend on model structure or model misspecification remains unknown. Future work should therefore include systematic sensitivity analyses across stopping criteria, prior families, noise levels, parameter selection, and alternative mechanistic models. Model-ensemble or model-averaged experimental-design approaches could also be considered so that a measurement schedule is selected based on its informativeness across several plausible descriptions of treatment response, rather than being optimal only for one assumed model. Such analyses would help identify scan schedules that remain stable across uncertainty scenarios and support the development of more robust data-collection plans for prospective use.

Another limitation is that the current implementation bases scan selection primarily on information represented by the simplified mechanistic model and therefore does not capture the full range of clinical knowledge used in practice. Clinician judgment may identify important features—such as atypical response patterns, comorbidities, treatment toxicity, or concern for rapid progression—that justify earlier or additional measurements even when the model predicts limited information gain. Future versions of INFORME could incorporate such expertise through clinician-informed priors, feasibility constraints, or additional terms in the design score, while preserving the ability to override an algorithmic recommendation when clinically necessary. The NLME component could also be extended to include richer patient-level covariates, including age, prior treatment history, baseline disease burden, comorbidities, imaging characteristics, and genomic features. These variables could inform individualized parameter priors, response-group assignment, and candidate scan schedules beyond what can be inferred from tumor volume measurements alone. Incorporating these complementary sources of information may improve personalization and clinical relevance, although careful validation will be required to avoid unstable subgrouping or overfitting when cohort sizes are limited.

Practical implementation of our framework must also account for operational constraints that are not represented in the current design formulation. MRI and other imaging appointments often need to be reserved well in advance, and the recommended scan time may not be available because of scanner capacity, staffing, treatment schedules, or patient-related factors. In addition, a scan planned for a specific day may be shifted because of illness, transportation difficulties, or other clinical needs. The present framework treats candidate measurement times as precisely available, whereas prospective use may require greater flexibility. Future extensions could incorporate scheduling availability, acquisition cost, and allowable timing windows directly into the design objective, producing a recommended interval or a ranked set of alternative dates rather than a single exact time point. The robustness of the resulting predictions should also be evaluated under realistic timing perturbations, delayed or missed scans, and substitutions with nearby feasible dates. Such modifications would help ensure that adaptive schedules remain informative even when the mathematically optimal scan time cannot be implemented exactly and would improve the framework’s practicality in clinical workflows.

Taken together, these results show that when a calibrated cohort is available, combining its hierarchical parameter distributions with an information-theoretic design criterion can identify measurement times that complete patient-specific predictions earlier and with fewer acquisitions than a fixed schedule. Prospective validation will require denser preclinical data, comparison with optimized fixed designs, and a design score that does not depend on the measurement being scheduled. The retrospective results provide a basis for this next step.

## Supporting information

### Data Availability

The source code is available on GitHub at https://github.com/heyrim/INFORME scan scheduling under the MIT License upon submission.

## Acknowledgments

We also acknowledge the Institute for Computational and Experimental Research in Mathematics (ICERM) at Brown University for supporting our participation in the *Uncertainty Quantification for Mathematical Biology* workshop, which provided an opportunity for discussions that helped shape this work.

## 1 Supplementary Material

### 1.1 Model fit to the synthetic prostate cancer cohort (HD1)

We calibrated the low-fidelity model for HD1 introduced in Methods to the 120 simulated tumor volume trajectories (3 groups) using NLME. Random effects were assigned to the intrinsic growth rate *r*, the carrying capacity *K*, and the treatment parameter *γ*_0_, each assigned a log-normal distribution. The response group was included as a categorical covariate on *γ*_0_ only, so that *r* and *K* share a single population distribution across groups while the treatment response remains group-specific. As for HD2, we present only the best variant tested here, since this work is not intended as a model selection procedure.

Table 2 summarizes the estimated population parameters. Including the response-group covariate on *γ*_0_ reduces the estimated inter-individual variability of that parameter from 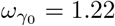 to 0.41, while leaving the estimates for *r* and *K* essentially unchanged (*r* = 0.32 with *ω*_*r*_ changing from 0.12 to 0.13; *K* = 0.57 with *ω*_*K*_ changing from 0.13 to 0.15). Most of the between-individual variability in treatment response is therefore accounted for by group membership, which is what makes the group-specific distributions informative as priors for the adaptive schedule. The estimated typical values of *γ*_0_ differ by roughly an order of magnitude between high and low responders. The random effects of *r* and *K* are negatively correlated, consistent with the well-known trade-off between growth rate and carrying capacity in logistic models. The sampling procedure uses marginal prior for each parameter, but an extension to incorporate the joint probability is straightforward. Moreover, all population parameters are precisely determined (R.S.E. ¡ 12%). Figs 6–8 show the individual fits across the cohort.

**Table 2.** Population parameter estimates for the model fit to HD1 (Monolix). S.E.: standard error (stochastic approximation). R.S.E.: relative standard error. C.V.: coefficient of variation, reported only for the random-effect standard deviations. P2.5 and P97.5 are the 2.5th and 97.5th percentiles of the estimate. Response group enters as a categorical covariate on *γ*_0_, with Group 1 (high responders) as the reference category; the fixed effects by category are obtained from *γ*_0,pop_ and the covariate coefficients 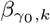 on the log scale, so *γ*_0,1_ coincides with *γ*_0,pop_. corr(*η*_*K*_, *η*_*r*_) is the estimated correlation between the random effects of *K* and *r*.

| Parameter | Value | S.E. | R.S.E. (%) | C.V. (%) | [P2.5, P97.5] |
| --- | --- | --- | --- | --- | --- |
| *Fixed effects* |  |  |  |  |  |
| $r_{\text{pop}}$ | 0.32 | 0.0038 | 1.20 | — | [0.31, 0.32] |
| $K_{\text{pop}}$ | 0.57 | 0.0079 | 1.39 | — | [0.56, 0.59] |
| $\gamma_{0,\text{pop}}$ | 4.12 | 0.27 | 6.56 | — | [3.62, 4.68] |
| $\beta_{\gamma_0,2}$ | -0.82 | 0.092 | 11.3 | — | [-1, -0.64] |
| $\beta_{\gamma_0,3}$ | -2.37 | 0.1 | 4.22 | — | [-2.57, -2.17] |
| *Fixed effects by category* |  |  |  |  |  |
| $\gamma_{0,1}$ (high) | 4.12 | 0.27 | 6.56 | — | [3.62, 4.68] |
| $\gamma_{0,2}$ (medium) | 1.81 | 0.12 | 6.56 | — | [1.59, 2.06] |
| $\gamma_{0,3}$ (low) | 0.38 | 0.029 | 7.62 | — | [0.33, 0.45] |
| *Standard deviation of the random effects* |  |  |  |  |  |
| $\omega_r$ | 0.13 | 0.0091 | 7.05 | 13.01 | [0.11, 0.15] |
| $\omega_K$ | 0.15 | 0.011 | 7.38 | 14.83 | [0.13, 0.17] |
| $\omega_{\gamma_0}$ | 0.41 | 0.033 | 8.09 | 43.10 | [0.35, 0.48] |
| *Correlations* |  |  |  |  |  |
| $\text{corr}(\eta_K, \eta_r)$ | -0.85 | 0.032 | 3.81 | — | [-0.9, -0.77] |
| *Error model parameters* |  |  |  |  |  |
| $a$ | 0.017 | 0.00015 | 0.877 | — | [0.017, 0.017] |

**Fig 6.**
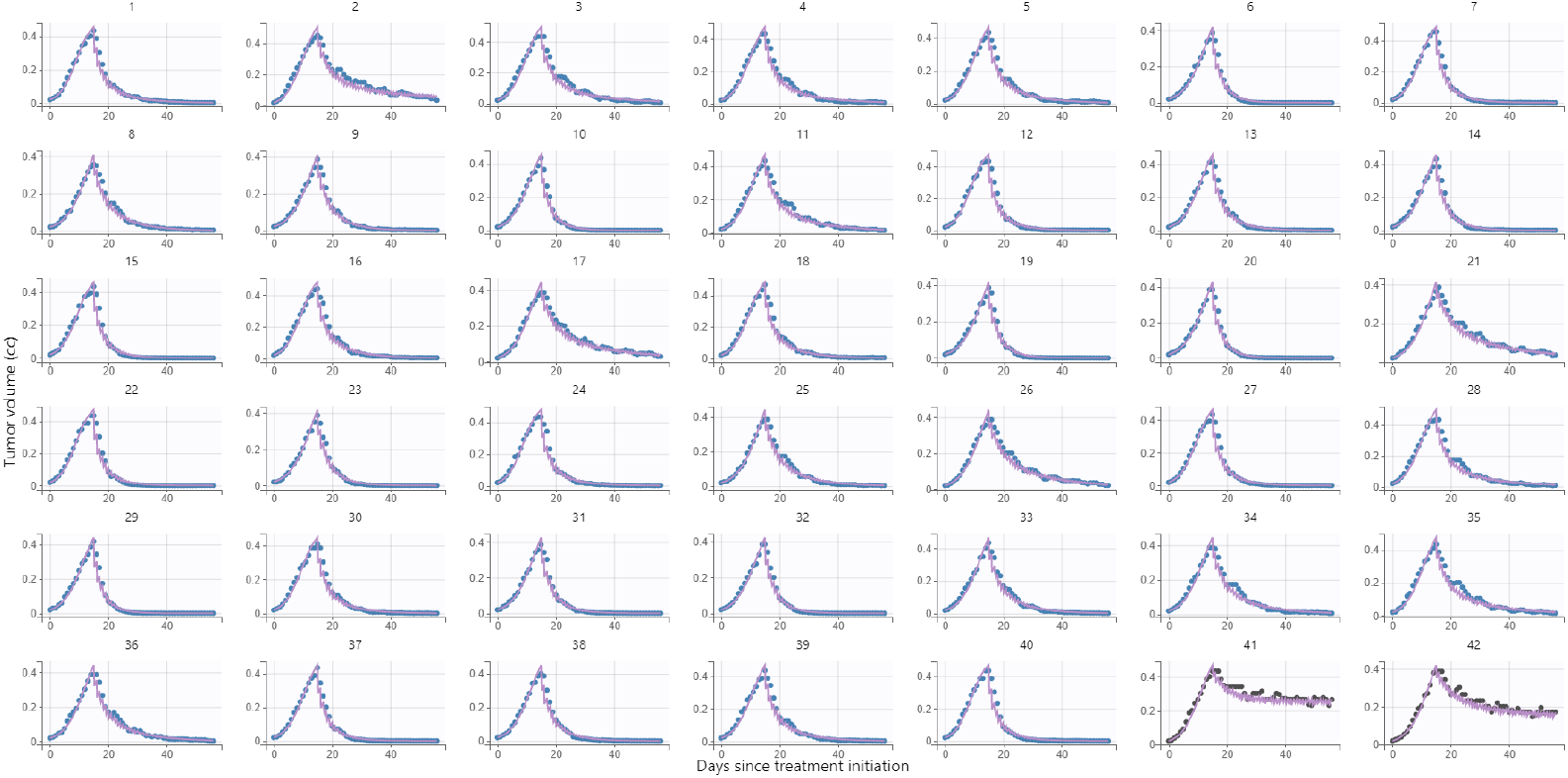
Individual model fits for HD1 (part 1 of 3). Colors show the response level: high responder (Group 1, blue), medium responder (Group 2, black), and low responder (Group 3, green).

**Fig 7.**
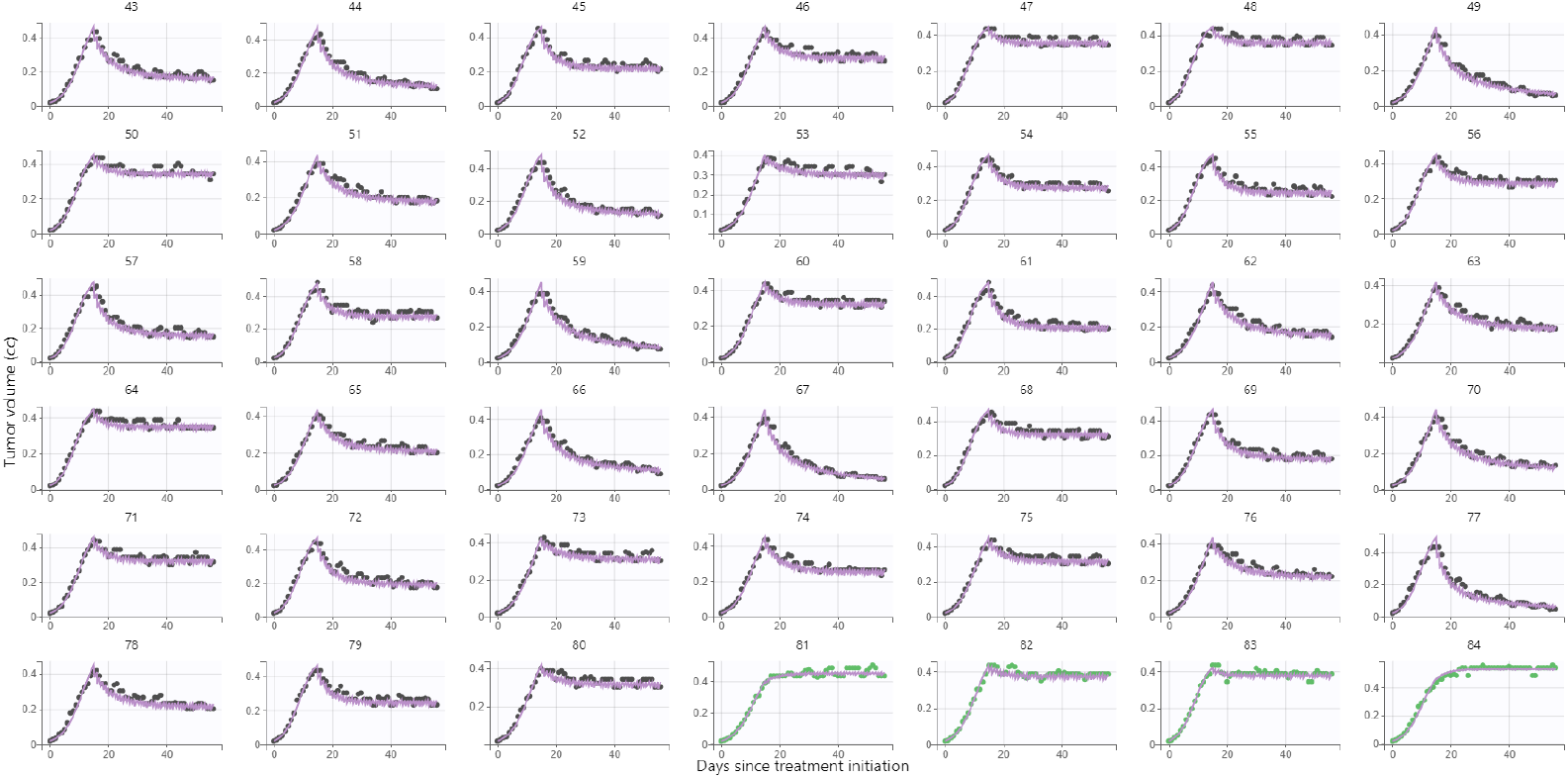
Individual model fits for HD1 (part 2 of 3). Colors show the response level: high responder (Group 1, blue), medium responder (Group 2, black), and low responder (Group 3, green).

**Fig 8.**
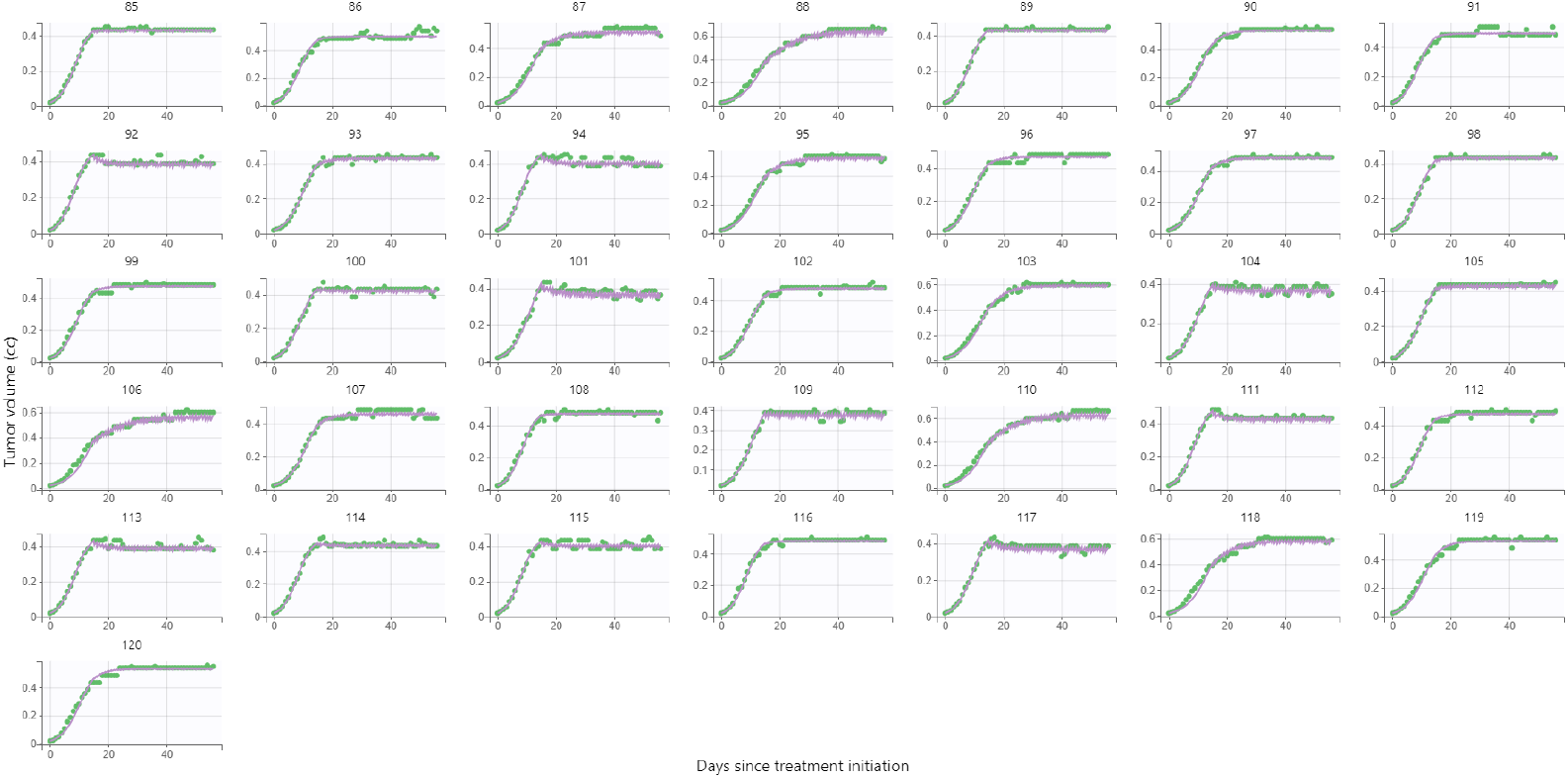
Individual model fits for HD1 (part 3 of 3). Colors show the response level: high responder (Group 1, blue), medium responder (Group 2, black), and low responder (Group 3, green).

### 1.2 Model fit to the head and neck cancer cohort (HD2)

For the model fitting, we assume a radiotherapy dose of 2 Gy per fraction, following Zahid et al. [32]. After treatment initiation (*t* = 0), treatment is administered on five consecutive days (Monday-Friday), followed by a two-day break (weekend), and this schedule is repeated until the last time point. Individual fits obtained from Monolix can sometimes appear poor because the algorithm constrains parameter variability across the population. The parameter fitting results and ranges used by Zahid et al. and the ranges used for sensitivity analysis by Mohsin et al. are different. For *λ*, we fix *λ* = 0.33 per day (or 0.01375 per hour) to reduce model identifiability issues. For PSI, we extend the range slightly from (0.1, 0.95) to (0.1, 1). For *δ*, we also extended the range followed by Zahid et al., which is (0, 0.1) to (0, 0.15). *V*_0_ is taken as the last tumor volume measurement before treatment initiation. Table 3 summarizes the parameter ranges. Table 4 summarize the estimated parameter values. Since all estimated parameters are well constrained, it suggests Monolix was able to estimate them robustly. Finally, model best fit is presented in Fig 9, which shows the simple model recapitulates data. We present only the best variant tested here, since this work is not intended as a model selection procedure.

**Table 3.** Parameter definitions, units, ranges and sources for Zahid et al. and Mohsin et al.

| Parameter | Definition | Unit | Range | Reference |
| --- | --- | --- | --- | --- |
| $\lambda$ | Intrinsic growth rate | $\text{h}^{-1}$ | 0.01375 (fixed) | Mohsin et al. [35] |
| PSI | $V(0)/K$ | unitless | (0.1, 1) | Mohsin et al. [35] |
| $\delta$ | $K$-reduction rate | unitless | (0, 0.15) | Zahid et al. [32] |

**Table 4.** Population parameter estimates for Model fit to HD2 (Monolix). S.E.: standard error (stochastic approximation). R.S.E.: relative standard error. C.V.: coefficient of variation, reported only for the random-effect standard deviations. P2.5 and P97.5 are the 2.5th and 97.5th percentiles of the estimate. *λ* was fixed (0.01375 per hour) and therefore has no associated uncertainty. The fixed effects by category are derived from *δ*_pop_ and the covariate coefficients *β*_*δ*,*k*_, so *δ*_1_ coincides with *δ*_pop_.

| Parameter | Value | S.E. | R.S.E. (%) | C.V. (%) | [P2.5, P97.5] |
| --- | --- | --- | --- | --- | --- |
| *Fixed effects* |  |  |  |  |  |
| $\lambda_{\text{pop}}$ | 0.014 | — | — | — | — |
| $\text{PSI}_{\text{pop}}$ | 0.87 | 0.022 | 2.54 | — | [0.82, 0.9] |
| $\delta_{\text{pop}}$ | 0.0058 | 0.0012 | 20.4 | — | [0.0039, 0.0085] |
| $\beta_{\delta,2}$ | 0.94 | 0.28 | 29.5 | — | [0.4, 1.49] |
| $\beta_{\delta,3}$ | 2.33 | 0.29 | 12.6 | — | [1.75, 2.9] |
| *Fixed effects by category* |  |  |  |  |  |
| $\delta_1$ | 0.0058 | 0.0012 | 20.4 | — | [0.0039, 0.0085] |
| $\delta_2$ | 0.014 | 0.0024 | 17.1 | — | [0.01, 0.019] |
| $\delta_3$ | 0.044 | 0.0066 | 15.0 | — | [0.032, 0.058] |
| *Standard deviation of the random effects* |  |  |  |  |  |
| $\omega_{\text{PSI}}$ | 0.72 | 0.14 | 19.9 | 10.49 | [0.49, 1.05] |
| $\omega_{\delta}$ | 0.47 | 0.092 | 19.7 | 46.51 | [0.32, 0.68] |
| *Error model parameters* |  |  |  |  |  |
| $a$ | 1.71 | 0.077 | 4.48 | — | [1.57, 1.87] |

**Fig 9.**
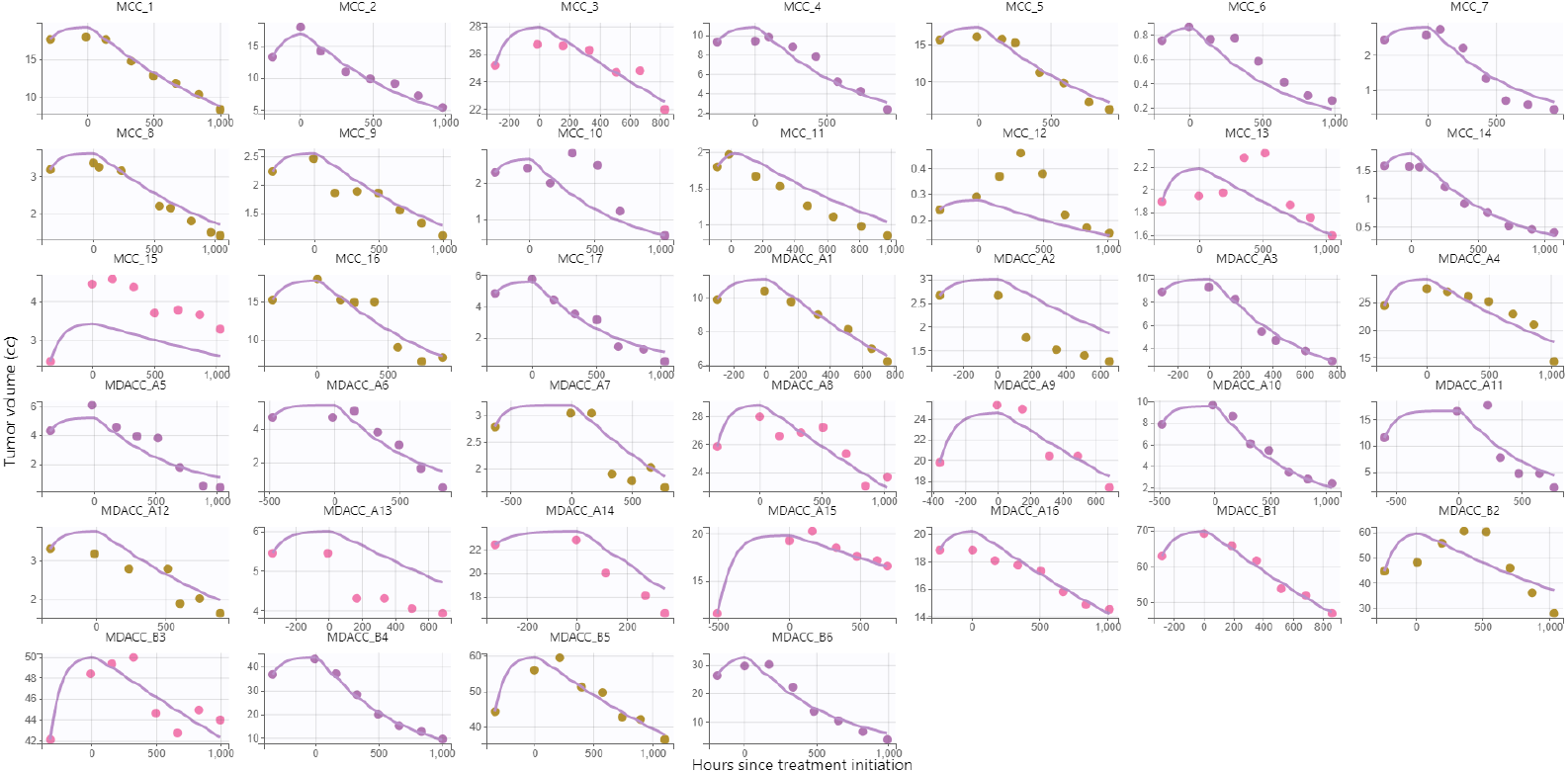
Model 7 with a covariate on *δ* based on response level. colors show the response level: high responder (purple), medium responder (brown), and low responder (pink).

## Notes

### Competing Interest Statement

The authors have declared no competing interest.

